# Estimating the Dynamic Range of Quantitative Single-Molecule Localization Microscopy

**DOI:** 10.1101/2021.05.24.445502

**Authors:** Daniel F. Nino, Joshua N. Milstein

## Abstract

In recent years, there have been significant advances in quantifying molecule copy number and protein stoichiometry with single-molecule localization microscopy (SMLM). However, as the density of fluorophores per diffraction-limited spot increases, distinguishing between detection events from different fluorophores becomes progressively more difficult, affecting the accuracy of such measurements. Although essential to the design of quantitative experiments, the dynamic range of SMLM counting techniques has not yet been studied in detail. Here we provide a working definition of the dynamic range for quantitative SMLM in terms of the relative number of missed localizations or blinks, and explore the photophysical and experimental parameters that affect it. We begin with a simple two-state model of blinking fluorophores, then extend the model to incorporate photobleaching and temporal binning by the detection camera. From these models, we first show that our estimates of the dynamic range agree with realistic simulations of the photoswitching. We find that the dynamic range scales inversely with the duty cycle when counting both blinks and localizations. Finally, we validate our theoretical approach on dSTORM datasets of photo-switching Alexa647 dyes. Our results should help guide researchers in designing and implementing SMLM-based molecular counting experiments.

## INTRODUCTION

Advances in fluorescence microscopy continue to push the limits of our knowledge of cellular and molecular biology. In particular, with roughly 25 years of steady improvements to super-resolution fluorescence microscopy, light microscopy is now able to reveal many of the finest details of the microscopic world [1–4]. Conventional fluorescence microscopy is fundamentally hindered by diffraction, limiting the resolution to hundreds of nanometers [5]. However, a resolution of about 10 nm is typically attainable with a subset of super-resolved methods that we refer to collectively as single-molecule localization microscopy (SMLM). SMLM includes techniques such as (d)STORM [6–8], (f)PALM [9, 10], and DNA PAINT [11].

While qualitative insight into organization and function is developed through high-resolution images, single-molecule imaging is rapidly evolving into a quantitative technique. Recently, significant effort has gone into quantifying molecular abundance and stoichiometry from SMLM imaging data [12–26]. These molecular counting methods seek primarily to correct for under- and over-counting artifacts introduced by multiple reversible transitions between the ON and OFF states used to achieve the sparsity conditions required for super-resolved imaging. The experimental parameters needed to extract molecule counts are then naturally related to the emission properties of the fluorophores, such as the number of localization events [14, 16, 22, 27], the number of blinks [13, 15, 17, 19, 21, 24], the ON and OFF times or rates [20, 25] –or some combination of these quantities.

Critical to any analysis is the need to correctly identify single emission events as detection errors affect the measured values of all the parameters just enumerated. As the density of fluorophores within a diffraction-limited volume increases, eventually, signal overlap will hinder the quantification of a molecule count. This limit, essentially, sets the dynamic range for molecular counting by SMLM. For diffraction-limited counting techniques such as stepwise photobleaching and counting by photon statistics, the dynamic range is well quantified. For example, the dynamic range of stepwise photobleaching is about 10 molecules per diffraction-limited spot [28], though recent innovations in the step-identification analysis have aimed to increase this number significantly [29, 30]. Likewise, counting by photon statistics currently has a dynamic range of about 25 fluorophores per diffraction-limited spot, limited mostly by the dead-time of the detection electronics [31].

In spite of the increasing number of publications on SMLM-based counting techniques, a detailed study of the dynamic range of these methods is lacking. While a handful of groups have briefly explored this topic, their efforts have been limited to a consideration of effects on image reconstruction, which is a more forgiving application than molecular counting. For example, early studies on the fluorophore densities needed for high structural resolution in image reconstructions pointed to a photoswitching duty cycle (*DC*) dependance that scaled inversely with the fluorophore density *DC* ≲ 1*/N* [32]. A more detailed estimate of the degree of overlap in detection events was published for qPAINT, but its effects were again limited to artifacts in image reconstruction [33]. An early estimate of the dynamic range for molecular counting was given by Annibale et al. [12]. They estimated a dynamic range of ∼ 1000 dyes*/µ*m^2^ through simulations for a single set of photokinetic parameters, but did not go further in their study or corroborate this experimentally. We previously estimated that SMLM counting techniques would have a dynamic range on the order of just a few hundred fluorophores per diffraction-limited spot [34]. However, this did not take into account effects due to photobleaching or the discretization of emissions into frames in the acquisition.

In this manuscript, we explore the dynamic range of SMLM-based molecular counting in detail through theoretical models and numerical simulations, which can be used to estimate the dynamic range given a set of photokinetic parameters and acquisition times. Our estimates are then experimentally validated using dSTORM data of Alexa647 on dsDNA templates, the results of which can be readily generalized to other SMLM techniques.

## MATERIALS AND METHODS

### Quantifying the dynamic range

A central quantity in estimating the dynamic range of SMLM counting techniques is the duty cycle (DC), typically defined as

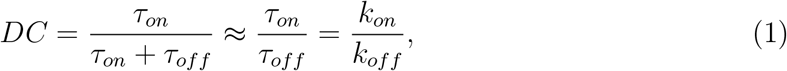

where *τ*_*on*_ is the average ON time of a fluorophore emission event, which we will refer to as a blink, and *τ*_*off*_ is the average OFF time. Likewise, *k*_*on*_ and *k*_*off*_ are the associated rates for switching into the ON and OFF states, respectively. To maintain sparsity *τ*_*off*_ ≫ *τ*_*on*_ or *k*_*off*_ ≫ *k*_*on*_. Under the assumption of negligible photobleaching, the duty cycle gives the probability of a fluorophore being in the ON state at some point in time. If one of *N* fluorophores within a diffraction limited volume is ON at time *t*, the probability that one of the other fluorophores is also ON would be given by (*N* − 1)*DC*, implying a dynamic range on the order of 1*/DC*.

A more quantitative way to asses the dynamic range of an SMLM measurement would be to consider how many localizations, from *N* fluorphores within a diffraction limited volume, one is expected to miss during the course of an experiment of total duration *T*. This may be expressed as follows:

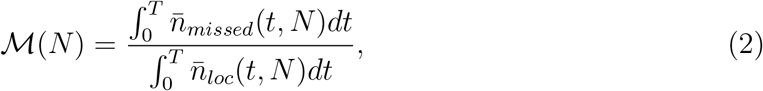

where 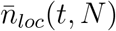 is the average number of localizations at time *t* and 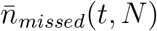 is the mean number of localizations we expect to miss due to signal overlap at time *t*. We first evaluate Eq. 2 for the case of negligible photobleaching, then move on to account for photobleaching effects.

### Two-state system

In the limit of negligible photobleaching, we consider a simplified two-state photoblinking model in which fluorophores can cycle indefinitely between an ON state and an OFF state with rates *k*_*on*_ and *k*_*off*_. The full dynamics of the probability of finding a fluorophore in the ON or OFF state can be obtained by solving a system of coupled ordinary differential equations describing the transitions (Supporting Material). If we assume that all fluorophores are OFF at the beginning of the acquisition (*t* = 0), then the probability of having a fluorophore ON at some later time *t* is given by:

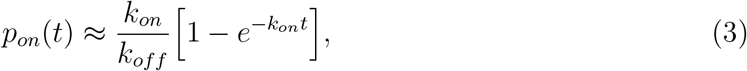

which holds so long as *DC* ≪ 1 or *k*_*off*_ ≫ *k*_*on*_.

Since we are interested in the probability of overlapping events, we need to consider the probability of having *n* fluorophores ON at the same time *t* within a diffraction limited volume, which we denote by *P* (*n*; *t*). If the emission of each fluorophore is independent of the other fluorophores, *P* (*n*; *t*) is simply a binomial distribution:

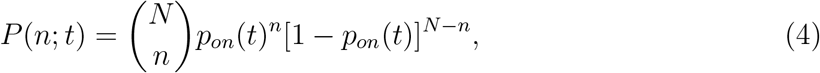

where *p*_*on*_(*t*) is given by Eq. 3 and *N* is the total number of fluorophores. The expected number of localizations at any one point in time *t* is then the expectation value

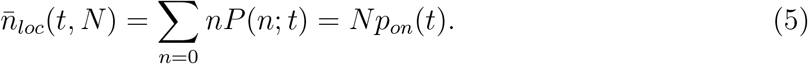

Now, if a subset of the fluorophores are all ON at the same time, we will underestimate the total number of localizations, effectively missing all but one of them. The expected number of missed localizations at time *t*, therefore, can be expressed as follows:

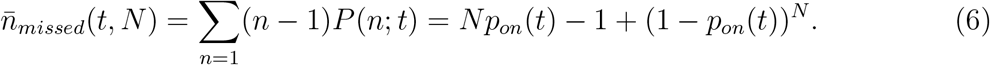

Transients in the photoblinking dynamics are generally short-lived relative to the total acquisition time. For instance, a typical SMLM acquisition will extend for 2 × 10^4^ frames each with a 50 ms exposure time, and with *k*_*off*_ ∼ 10^−2^ ms^−1^ and *k*_*on*_ ∼ 10^−5^ ms^−1^ as typical photokinetic rates [7, 13, 35]. The photoblinking dynamics, therefore, quickly reaches the steady-state in which the probability of finding a fluorophore ON at any one time step simplifies to 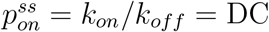. At steady-state, the integrals in Eq. 2 can be evaluated and we arrive at the relation

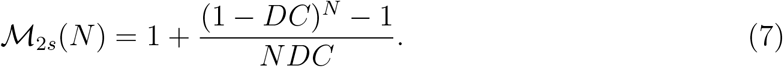

So long as *DC* ≪ 1 and *NDC* ≪ 1, this may be approximated as

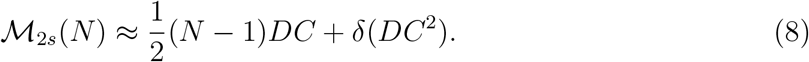

### Three-state system

In spite of improved imaging buffers that mitigate the effects of photobleaching, all fluorophores used in SMLM will eventually, irreversibly photobleach. The simplest way to incorporate photobleaching is to include a third state in the model that allows an irreversible transition from the ON state to the photobleached (PB) state with rate *k*_*pb*_. From the resulting system of differential equations (Supporting Material), the probability of having a fluorophore ON at some time *t* is given by:

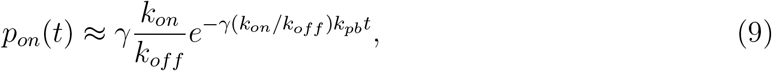

where *γ* = 1*/*(1+*k*_*pb*_*/k*_*off*_) and we have assumed that *k*_*pb*_ ∼ *k*_*off*_ ≫ *k*_*on*_. The relative number of missed localizations *ℳ*_3*s*_(*N*), Eq. 2, must now be evaluated numerically. However, a lowest order analytic approximation, similar to that found for the two-state model in Eq. 8, is given by the following:

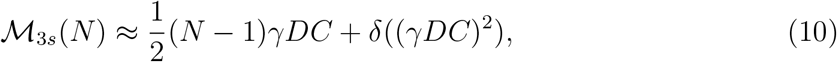

assuming *γDC* ≪ 1, *NγDC* ≪ 1, and *T* → *∞* for this three-state model.

### Simulations in ‘continuous’ and ‘discrete’ time

To simulate blinking fluorophores, we assume Markovian dynamics for the transitions between the different states available to each fluorophore. To ensure first order kinetics, the time step is taken to be *δt* = 1 ms, much smaller than the time scale of any of the possible transitions. We refer to these simulations as ‘continuous’ time simulations to differentiate from those that account for temporal binning by the detection camera, which we will refer to as ‘discrete’ time simulations. The probability of transition between any state *i* and *j* is assumed to follow first-order kinetics, i.e. Prob(*i* → *j*) = *k*_*i,j*_*δt* + *O*(*δt*^2^) where *k*_*i,j*_ is the transition rate between state *i* and state *j*, and we ignore contributions of order *O*(*δt*^2^). The emission of *N*_*total*_ dyes over *T* time steps is encoded in a *N*_*total*_ × *T* matrix Σ. If dye *N*_*l*_ is ON at time step *t*_*i*_, then the matrix element (*N*_*l*_, *t*_*i*_) of Σ is 1. Conversely, if dye *N*_*l*_ is OFF or photobleached at time step *t*_*i*_, then the matrix element (*N*_*l*_, *t*_*i*_) of Σ is 0. Each row in this matrix then is a simplified representation of the emission time series for a single fluorophore that ignores intensity fluctuations.

To obtain a numerical measure of the dynamic range for a given number of fluorophores *N*, a set of *N* rows is randomly chosen from Σ without replacement so as to not choose the same fluorophore in any one sample. The total number of localizations and blinks from the distinct fluorophores chosen in this process is then recorded. The selected rows are then added together, mimicking the case for *N* fluorophores within a diffraction-limited spot. From the resulting combined time series, the total number of localizations and blinks are recorded yielding a measure of the total number of missed events. To augment the number of samples, we bootstrap across different combinations of *N* distinct rows to obtain the average relative number of missed events *ℳ*.

Actual single-molecule localization microscopy acquisitions are taken over discrete frames, typically several tens of milliseconds long. This ensures the collection of enough photons to obtain a high localization precision. To account for this, we discretize the continuous time simulations into frames of duration *τ*_frame_. This creates a new matrix Σ′ containing the discretized emission time series of each fluorophore. If, over the course of a frame, a fluorophore is ON for at least one time step within the frame, it is also considered ON for the entire frame and the value of that element in Σ′ is taken to be 1. If the fluorophore is not ON in any time step within a frame, the corresponding entry in Σ′ is 0.

### Chambers and surface functionalization

Chambers were made by drilling holes into glass slides with diamond powder coated drill bits. To create flow chambers, 300LSE double-sided adhesive film (3M, Saint Paul, USA) was cut using a plotter cutter (CM350e, Brother, Dollard-des-Ormeaux, Québec) in the shape of rectangular flow channels. This was used to attach a cover slip (VWR micro cover glass, 22×50 mm No 1.5, VWR International, Mississauga, Ontario) to the slides so as to form 4 parallel, independent flow chambers with 2 fluid access ports for each chamber.

Surface functionalization was performed as in *Nino et al*. [34]. In brief, cover slips were incubated in 1% v/v poly-l-lysine solution overnight. A 5 % (v/v) solution of gold nanoparticles (40 nm diameter, Sigma Aldrich, St. Louis, Missouri) was flowed into a given chamber and incubated for 15 minutes. The flow was then flushed with 100 *µ*L of distilled water. Sparse gold nanoparticles bound to the surface were used as fiducial markers in post-processing for lateral drift correction. The surface of the flow chamber was then functionalized by flowing 50 *µ*L of 0.5 mg/mL BSA-Biotin. After a 10 minute incubation, the chamber was flushed with 100 *µ*L of PBS (pH 7.4, filtered through a 0.2 *µ*m filter). To further block the surface, 50 *µ*L of 10% (w/v) BSA solution was added into the chamber and incubated for 5 minutes. After flushing with 100 *µ*L of PBS, 50 *µ*L of 0.5 mg/mL of streptavidin was flowed into the chamber and incubated for 10 minutes, after which point the chamber was flushed with 100 *µ*L of PBS. Finally, 20 bp biotinylated dsDNA, conjugated to a single Alexa647 at the opposite end, was incubated at 10 pM concentration for 10 minutes. This yielded a sparse distribution of single, surface-tethered fluorophores.

### dSTORM imaging

We employed continuous 2.4 kW*/*cm^2^ of 637 nm illumination and 12 W*/*cm^2^ of 405 nm illumination, the latter to drive fluorophores back to the ground singlet state from the long-lived dark state. We found that in the absence of 405 nm illumination, the OFF time distribution was not mono-exponential, likely indicative of a second dark state (see Fig. S3 in the Supporting Material.). The imaging buffer was the same for all experiments and consisted of 143 mM *β*-mercaptoethanol (BME), and the PCA/PCD oxygen scavenging system (13 mM and 50 nM, respectively). Further details of the microscopy instrumentation can be found in *Nino et al*. [34]. For each experiment, 25,000 frames were collected with an EMCCD camera (Andor iXon, Concord, MA). Each frame was 50 ms in duration, totalling approximately 20 minutes per acquisition with an electron-multiplication gain of 270. To generate a localization table, we used rapidSTORM [36] with a varying intensity threshold ranging from 8,000 to 10,000 photo-electron counts, using least-squares fitting with a fixed point-spread function (PSF) full-width at half maximum of 390 nm. The intensity threshold was chosen as low as possible so that both spurious, low intensity localizations and discarded true localizations were minimized. This threshold sometimes varied between acquisitions, so an intensity histogram was used to cutoff the intensity threshold at the point where the number of localizations began to sharply increase.

## RESULTS

### Continuous time simulations

To explore the dynamic range of SMLM-based molecular counting, we initially performed two sets of simulations in continuous time. We first considered 2000 fluorophores described by a two-state photoswitching model with an average ON time *τ*_*ON*_ = 100 ms and duty cycles ranging from *DC* = 10^−3^ to 10^−2^. We then considered 2000 fluorophores with the same ON and OFF rates as the two-state model, but which could also undergo photobleaching with rate *k*_*pb*_ = 5.3 × 10^−3^ ms^−1^. These photokinetic rates are equivalent to an average number of blinks per fluorophore of about 3, which is a typical value one might expect to observe for common fluorophores used in SMLM [37]. We use these simulations to verify Eq. 2 for the two-state and three-state models. The results are displayed in Fig. 1, which shows that the relative number of missed localizations for both the simulated two-state and three-state models agree with our theoretical estimates.

**FIG. 1.**
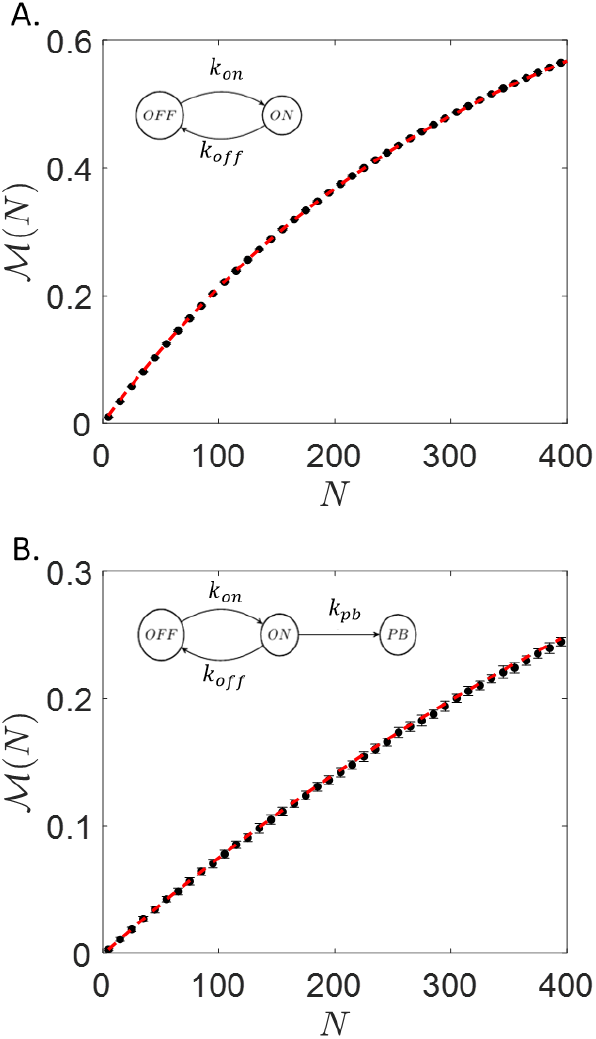
Relative number of missed localizations *ℳ* (*N*) as a function of the fluorophore density (*N* per diffraction limited volume) A.) without photobleaching and B.) with photobleaching (*k*_*pb*_ = 5.3 × 10^−3^ ms^−1^). For all simulations *k*_*on*_ = 5 × 10^−5^ ms^−1^ and *k*_*off*_ = 1 × 10^−2^ ms^−1^. Results from simulations with error bars estimated by bootstrapping (•). Theoretical estimates for the given photokinetic model (dashed lines).

Incidentally, the dynamic range improves when the fluorophores photobleach. Specifically, for higher photobleaching rates, higher fluorophore densities can be accommodated for a given relative number of missed localizations. The improvement scales linearly with an increasing photobleaching rate (see Fig. S1 in the Supporting Material). This implies that fluorophores that undergo less blinking cycles before photobleaching are better suited for applications requiring high labeling densities, as one would expect.

These results can be generalized to estimate the fluorophore densities that result in a certain relative number of missed localizations for a range of duty cycles. For each set of simulations, the average fluorophore density for which 5%, 10%, 15% and 20% of localizations were missed was plotted as a function of duty cycle (see Fig. 2). For each photokinetic model, we estimate the fluorophore densities by interpolating our simulation results and by a numerical evaluation of Eq. 2. We find that the fluorophore density, to reach a given relative number of missed localizations, scales inversely with the duty cycle (*N ∝* 1*/*DC) (insets) for both models. This agrees with the scaling previously reported in [32], but is now built upon a theoretical foundation from which exact estimates can be computed given a set of photokinetic parameters and including photobleaching effects.

**FIG. 2.**
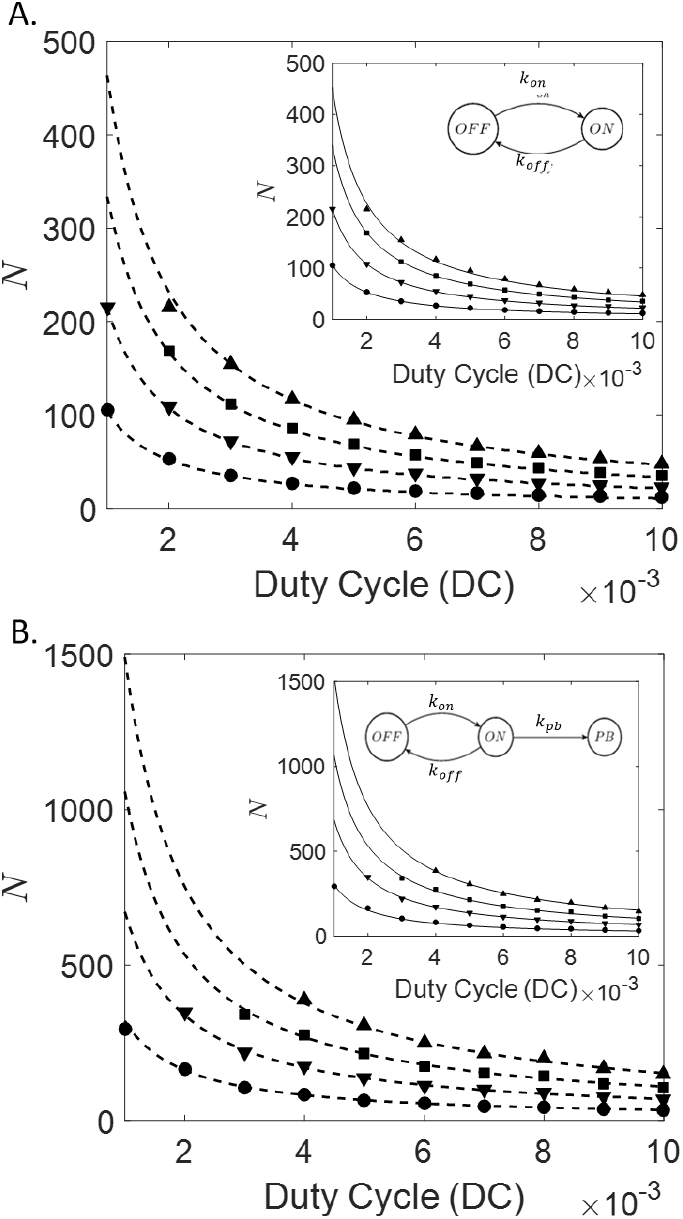
Simulated fluorophore densities (*N* per diffraction limited volume) for which 5%(•), 10%(▾), 15%(▪) and 20%(▴) of localizations are missed for the two-state (top) and three-state (bottom) models. Respective fluorophore densities estimated by Eq. 2 (dashed lines). Insets: Fluophore densities extracted from the simulations fit to *A/*DC (solid line), where *A* is a fitting constant.

### Accounting for the exposure time

While we have shown that our theoretical model holds rigorously in ‘continuous’ time (i.e. for small time steps compared to the time-scale of transitions between states), actual SMLM experiments are temporally limited by the camera acquisition rate. It can be expected that our estimates, so far, serve as an upper-bound on fluorophore densities given the loss of time resolution that results from acquiring frames tens of milliseconds long.

The expressions for the probability of a fluorophore being ON at any point in time (Equations (3) and (9)) need to account for the effects of an exposure time *τ*_frame_. Consider the probability of a dye being ON at some point during a frame. This can happen in two mutually exclusive ways: either the dye was already ON at the beginning of the frame, or the dye was OFF at the beginning of the frame and turned ON at some point during the frame (Supporting Material). These considerations lead to a scaling between the continuous time probabilities and the discrete time probabilities given by the linear relationship (see Fig. S2 in the Supporting Material):

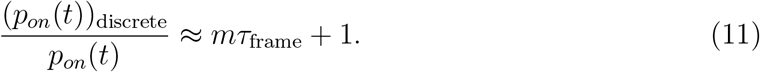

Here the slope is simply *m ≈ k*_*off*_ + *k*_*pb*_ for the three-state model and *m ≈ k*_*off*_, for the two-state model (Supporting Material). This expression indicates that the binomial model continues to hold for discrete time, provided that the probabilities are scaled according to Equation (11). Fig. 3 confirms this scaling for simulated data displaying the relative number of missed localizations, with and without the corrected scaling (*k*_*off*_ = 0.01 ms^−1^, DC = 5 × 10^−3^ and *k*_*pb*_ = 1.25 × 10^−3^ ms^−1^). To obtain error bars for the simulation results, the combined time series of *N* fluorophores were bootstrapped fifty times to obtain different combinations of overlapping fluorophores. For each sample of *N* fluorophores, the relative number of localizations were recorded. The mean and standard deviation of the 50 bootstrapped samples were used for each value of *N*. It should be noted that the linear approximation to the correction factor is quite accurate, even for exposure times much longer than the average ON time.

**FIG. 3.**
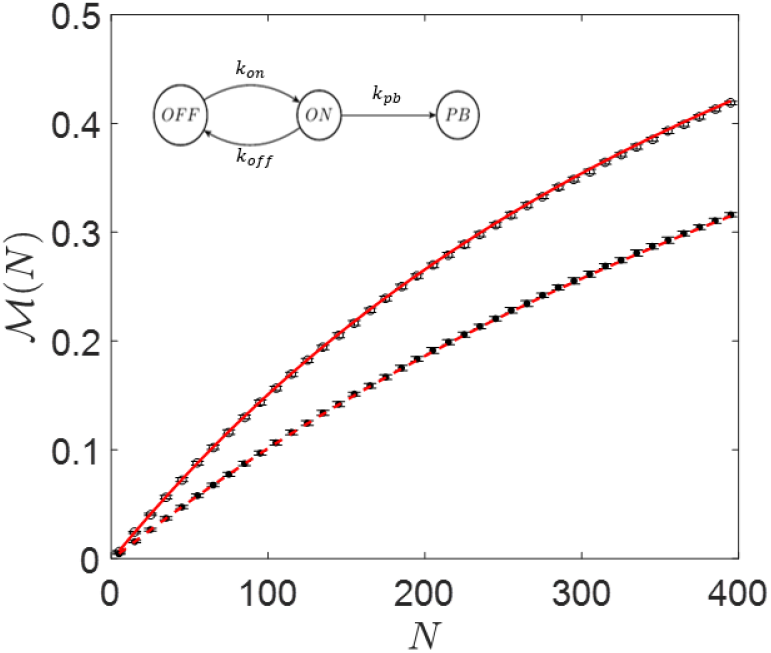
Exposure time effects on the relative number of missed localizations *M*(*N*) as a function of fluorophore density (*N* per diffraction limited volume). Simulations discretized into frames (∘) (*τ*_frame_ = 50 ms, *k*_*off*_ = 0.01 ms^−1^, DC = 5 × 10^−3^ and *k*_*pb*_ = 1.25 × 10^−3^ ms^−1^). The same simulation without temporal binning (•). The solid (dashed) curves show the theoretical estimates for discrete (continuous) time.

### SMLM counting experiments

Having developed a theoretical approach to estimating the relative number of missed localizations, which agrees with our simulations, we now validate it on experimentally acquired data. To mimic having an arbitrary, yet known, number of fluorophores within a diffraction-limited spot, we first acquire a stack of dSTORM images taken of single, Alexa647 dyes sparsely distributed on a coverslip. From the resulting localization table, we build a series of multiple-dye clusters by grouping the localizations of the individual dyes into diffraction limited spots. Of course, this approach ignores interactions between dyes that may take place in practice, but our theory also does not account for these effects at the moment.

The photokinetic rates were measured from the localization tables obtained for each acquisition. The distributions for the ON and OFF times were fit to an exponential function (Fig. 4A and 4B). The photobleaching rate was obtained by fitting the cumulative number of localizations to the cumulative distribution function of an exponential as shown in Fig. 4C (Supporting Material). Although a more careful analysis is needed to precisely extract the photokinetic rates [38], our inference of these rates from the localization table is sufficient for the present analysis. Finally, the relative number of missed localizations was measured for the experimentally acquired data sets and agreed well with our theoretical estimates. (Fig. 4D). Note, we have included error bars on the theoretical estimates because the underlying photokinetic parameters, which go into the model, must be measured. The error bars were estimated by sampling each parameter 100 times from a normal distribution with a corresponding mean 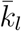 and standard deviation *δk*_*l*_, where *l* = ‘on’, ‘off’, or ‘pb’, obtained from the fits. From each sampled set of parameters, the relative number of missed localizations was calculated using Eq. 2 with one standard deviation within the distribution indicated by the error bars.

**FIG. 4.**
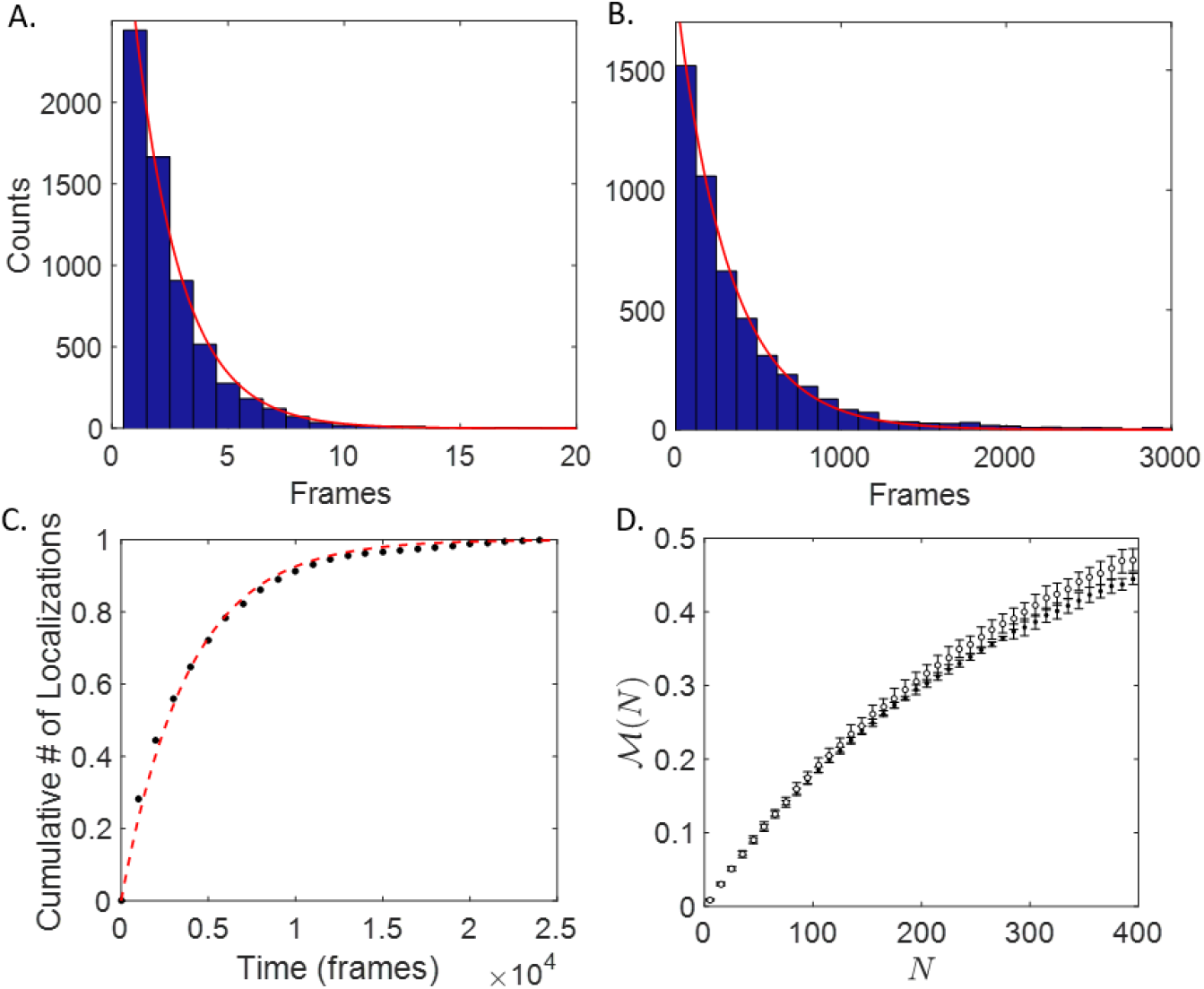
Experimental dSTORM data for photoswitching of Alexa647. A) Distribution of ON times fit to a single exponential (solid line) resulting in *τ*_*on*_ = 2.0 ± 0.1 frames. B) Distribution of OFF times fit to a single exponential (solid line) resulting in *τ*_*off*_ = 323 ± 3 frames. This yields a duty cycle of *DC* = (6.2 ± 0.3) × 10^−3^. C) Extracting the photobleaching rate using the normalized cumulative number of localizations. Experimental data (•). Fit to *f* (*t*) = 1 − *e*^−*ct*^ (dashed line), where *c* is the fitting constant and *t* is the time (in frames). The photobleaching rate *k*_*pb*_ = 0.044 ± 0.005 frame^−1^ is inferred from the fitting constant (Supporting Material). D) Experimental results for the relative number of missed localizations *ℳ* (*N*) as a function of fluorophore density (*N* per diffraction limited volume). Experimental data (•). Theoretical estimate (*∘*).

The slight discrepancy in Fig. 4D between the experimental data and the theoretical curve can likely be attributed to inaccuracies in our simple, three-state photokinetic model. Further analysis of the distribution of OFF times reveals a long tail which is better captured by fitting a double exponential model (see Fig. S3). This is likely a remnant of a second dark state whose transition appears to be suppressed by 405 nm irradiation. As previously mentioned, in the absence of 405 nm irradiation, the OFF time distribution is clearly modeled by a double exponential function, indicative of at least two dark states. With 405 nm irradiation, the OFF time distribution is fairly well captured by a single exponential fit. However, with such low 405 nm irradiation, kept low to keep the duty cycle low, the transition to the second dark state is not completely suppressed. While our model could in principle be generalized to account for this, it is difficult to correctly account for the possible transitions and to measure these rates accurately. For example, adding a transition from the long-lived dark state to a longer-lived dark state would then require a method to extract this transition rate experimentally as well as the transition rate from the second dark state back to the ground singlet state. While technically possible with hidden Markov models [39], it is not yet clear what the true pathways are for Alexa647 and more studies such as those performed in [40] need to be done.

### Extrapolating from missed localizations to missed blinks

So far, we have explored the dynamic range in terms of the relative number of missed localizations. However, for many SMLM-based counting techniques the dynamic range is actually set by the relative number of missed blinks per diffraction-limited spot, each of which may result in multiple, temporally adjacent localizations. Unfortunately, estimating missed blinks is more difficult than estimating missed localizations. However, we can estimate a lower bound under the assumption that the blinks are temporally sparse such that it is unlikely for more than two blinks to overlap at any moment in time. For a blink of duration *τ*_*on*_, which begins at time *t*′, a second blink of the same duration will overlap if it occurs between *t*′ − *τ*_*on*_ and *t*′ + *τ*_*on*_, resulting in a temporal overlap window of duration 2*τ*_*on*_ (see Fig. S4 in the Supporting Material). The factor of 2 translates to a scaling of *p*_*on*_(*t*) _Blinks_ *≈* 2 *p*_*on*_(*t*) _Locs_, which can then be substituted into our previous expressions to yield an analytic estimate of the relative number of missed blinks. Again, this simple scaling of *p*_*on*_ provides a lower bound on the relative number of missed blinks and will be most accurate at low fluorophore density.

Fig. 5A shows the results of simulations of a three-state model, indicating the number of fluorophores as a function of duty cycle that would result in missing 5%, 10%, 15% and 20% of the blinks. Of particular interest is that the dynamic range continues to scale inversely with the number of fluorophores (inset to Fig. 5A). It should be noted that, for a given duty cycle, the fluorophore density needed to obtain a certain relative number of missed blinks is about half that needed to obtain the same relative number of missed localizations. For example, for a duty cycle of 1× 10^−3^ in the two-state photoswitching model, on average, about 55 dyes result in 5% missed blinks, while about 105 dyes are need to miss 5% of the localizations. Fig. 5B shows that the theory agrees with the simulation results for the relative number of missed blinks as a function of fluorophore number up to about *ℳ*_*B*_ ∼ 0.2. For higher fluorophore densities, as expected, the theoretical estimate increasingly underestimates the actual relative number of missed blinks. Finally, Fig. 5C shows a similar level of accuracy by our theoretical approach at estimating the relative number of missed blinks when applied to our dSTORM data on Alexa647. Again, the error bars included in the theoretical estimate result from uncertainty in the model photokinetic parameters and are obtained through the bootstrapping procedure already described.

**FIG. 5.**
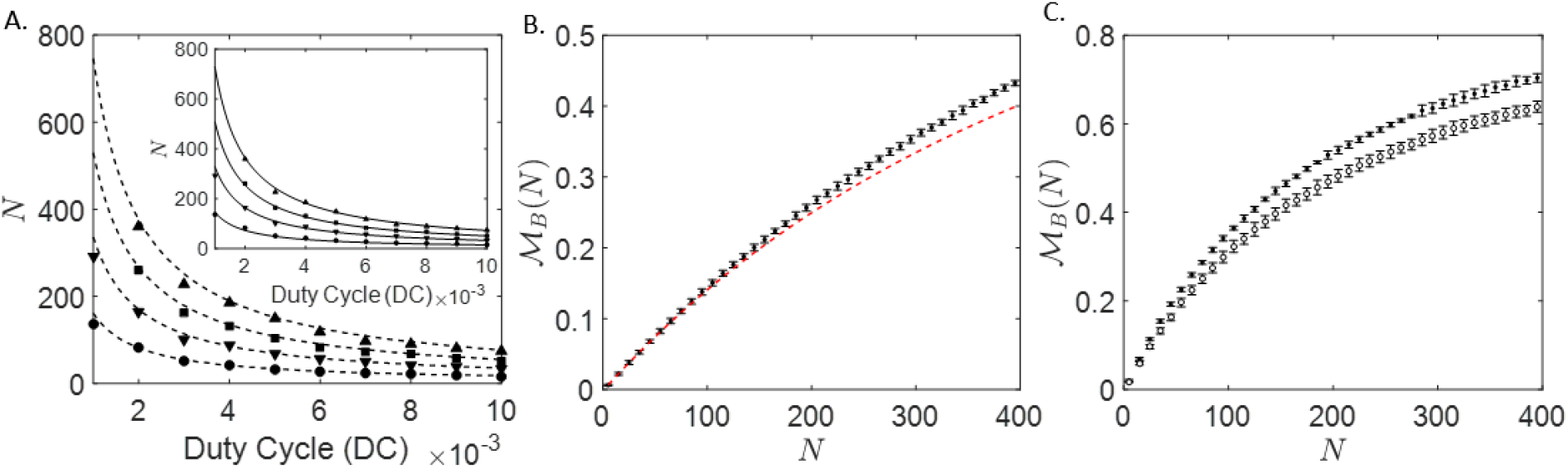
A) Fluorophore densities *N* as a function of duty cycle (*DC*) from simulations in continuous time. Markers indicate 5%(•), 10%(▾), 15%(▪) and 20%(▴) of the blinks are missed. Respective fluorophore densities estimated by Eq. 2 (dashed lines). Insets: Fluorophore densities extracted from the simulations fit to *A/*DC (solid line), where *A* is a fitting constant. B) Relative number of missed blinks *ℳ*_*B*_(*N*) as a function of fluorophore density *N* in continuous time (*k*_*off*_ = 0.01 ms^−1^, DC = 5 × 10^−3^ and *k*_*pb*_ = 5.3 × 10^−3^ ms^−1^). Simulated data (•). Theoretical estimate (dashed line). C) Relative number of missed blinks *ℳ*_*B*_(*N*) as a function of fluorophore density *N* for dSTORM data on Alexa647 (*k*_*on*_ = 0.50 ± 0.03 frame^−1^, *DC* = (6.2 ± 0.3) × 10^−3^, and *k*_*pb*_ = 0.044 ± 0.005 frame^−1^). Experimental data (•). Theoretical estimate (∘).

## DISCUSSION

In this manuscript, we have developed theoretical expressions for estimating the dynamic range of quantitative SMLM in terms of either the relative number of missed localizations or blinks. These expressions not only set a constraint on the number of molecules within a diffraction limited volume that can be counted, but clarify the relationship between the dynamic range and photophysical properties of the fluorescent labels, such as the photo-bleaching rate, duty cycle, etc., while accounting for a finite exposure time. Our results should both facilitate and improve the experimental design of quantitative SMLM measurements.

For instance, extremely low duty cycles in the range of *DC* ∼ 10^−4^ − 10^−5^ are commonly used for imaging and can be achieved with a variety of fluorophores [37, 41]. For similar photophysical parameters to those considered in our simulations, but with a *DC* ∼ 10^−4^ − 10^−5^, an extrapolation of Figure 2B implies that several 10^3^ fluorophores per diffraction limited volume could be counted by quantitative SMLM while missing less than 5% of the localizations. Dependent upon the required spatial precision, many SMLM-based counting applications might be able to employ a significantly higher duty cycle than this, while maintaining the necessary level of accuracy in a count. A higher duty cycle could be achieved by using a different fluorophore, or in the case of Alexa647, by using more intense 405 nm irradiation to rapidly induce the transition from the OFF to the ON state, effectively reducing the OFF time. For instance, in applications involving the quantification of protein stoichiometry, where much lower molecule densities can be expected, duty cycles of (*>* 10^−2^) could be used to expedite data collection and acquire better statistics. This would, of course, also depend upon the spatial resolution required to distinguish the individual macromolecular complexes under investigation, but for certain systems this may indeed be possible. The ability to increase the duty cycle to such an extent would reduce the total acquisition time and expand the range of systems where quantitative SMLM counting techniques may be applied, such as within slow-moving dynamical systems in live-cells.

## AUTHOR CONTRIBUTIONS

Nino and Milstein designed the research, developed the theory and carried out the simulations. Nino performed the experiments and analyzed the data. Nino and Milstein co-wrote the article.

## ACKNOWLEDGMENTS

DFN would like to thank Claudiu Gradinaru for fruitful discussions which inspired this study. This work was supported by a Natural Sciences and Engineering Research Council (NSERC) Discovery Grant and an Early Researcher Award from the Ontario Ministry of Research, Innovation and Science. DFN was supported by a MITACS Research Training Award.

## Supplementary Material

### 1 Two-state model

For a two-state model of blinking fluorophores, the probabilities *p*_*on*_(*t*) and *p*_*off*_ (*t*), for finding a fluorophore in either the ON or OFF state at some time *t*, are determined by the following system of coupled differential equations:

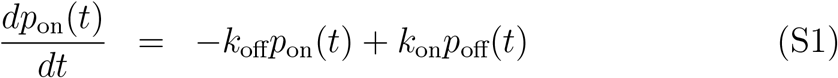

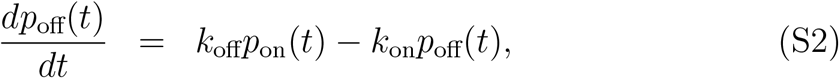

where *k*_off_ and *k*_on_ are the transition rates from the ON and OFF states, respectively. If all fluorophores are initially in the OFF state at time *t* = 0, these equations may be solved to yield

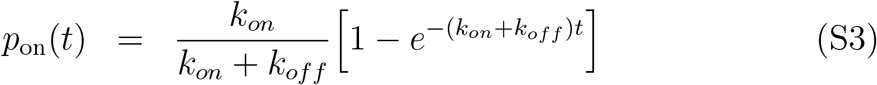

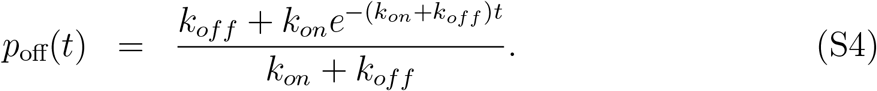

In steady-state (ss), we get the expected result of 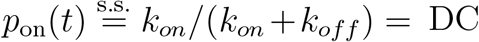 and 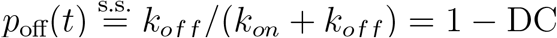.

### 2 Three-state model

To account for photobleaching effects, we add an irreversible transition from the ON state to a photobleached (PB) state. This results in the following set of coupled differential equations

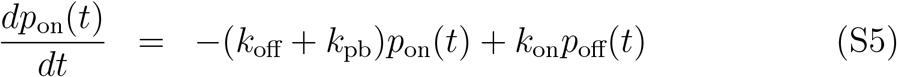

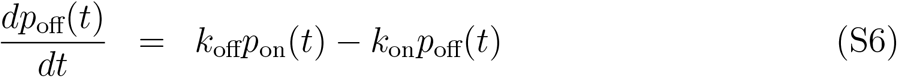

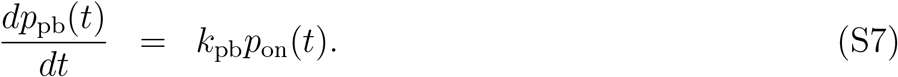

Once again, if all fluorophores are initially in the OFF state at *t* = 0, then solving for the probabilities yields the following:

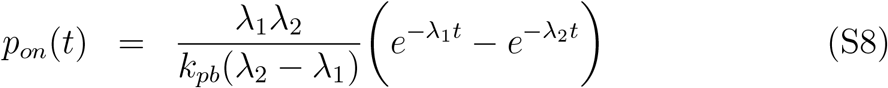

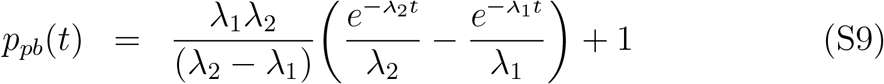

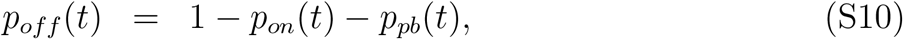

where *λ*_1_ = *k*_*on*_*k*_*pb*_*/k, λ*_2_ = *k*(1 − *k*_*on*_*k*_*pb*_*/k*^2^), and *k* = *k*_*on*_ + *k*_*off*_ + *k*_*pb*_. Under typically employed conditions for SMLM, *k*_*off*_ ∼ *k*_*pb*_ ≫ *k*_*on*_. This means that *λ*_2_ − *λ*_1_ *≈ λ*_2_ and that *λ*_2_ ≫ *λ*_1_, so 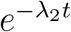 decays much faster than 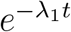 and we can take 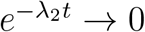. With these approximations

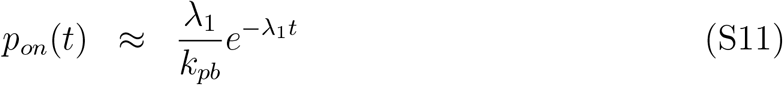

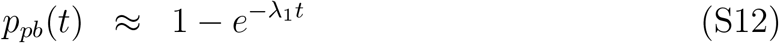

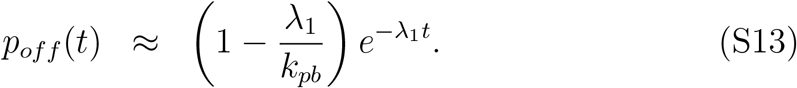

### 3 Measuring the photobleaching rate

The photobleaching rate can be measured experimentally by fitting the cumulative number of localizations as a function of time to a single exponential *F* (*t*) = 1 − *e*^−*ct*^. Since 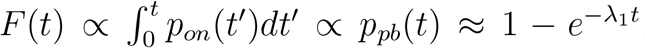, we can identify *c* = *λ*_1_. Isolating for the photobleaching rate, we are left with the following relation:

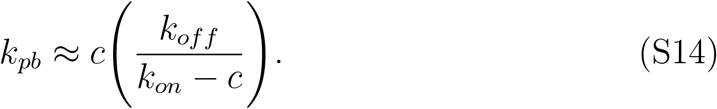

### 4 Effects of photobleaching

The average number of fluorophores needed to miss a given relative number of localizations and blinks scales linearly with the photobleaching rate. This points to the idea that fluorophores which photobleach faster (or equivalently blink less) are preferable for molecular counting applications.

**Figure S1:**
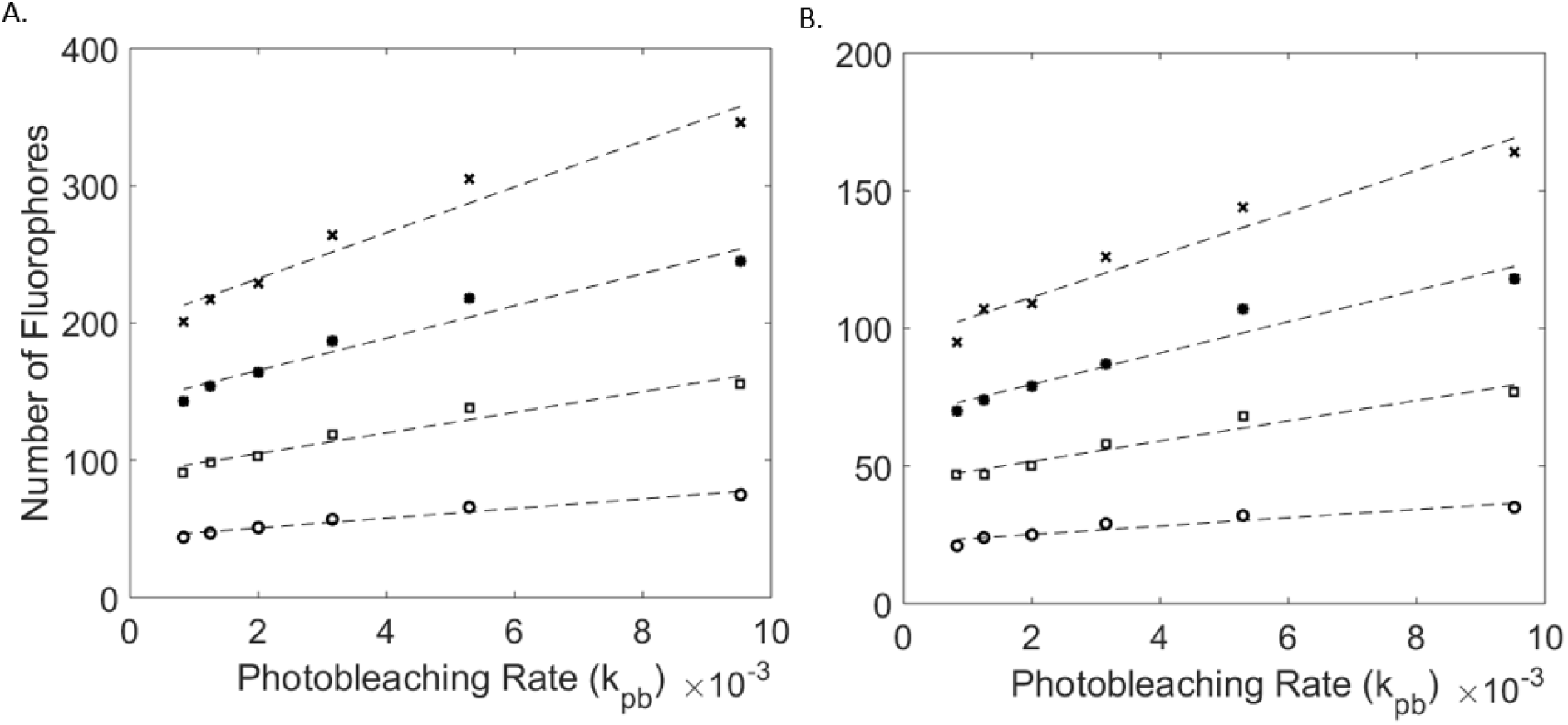
Average number of fluorophores for which 5% (circles), 10% (squares), 15% (stars), and 20% (crosses) of A) localizations and B) blinks are missed as a function of the photobleaching rate in the three-state photo-switching model.

### 5 Correcting for the exposure time

If a fluorphore is in the ON state at any moment during a single camera frame, we will assume that the camera is able to detect the emission, resulting in a single localization for that frame. We will also assume there are two ways to achieve a localization event, either the fluorophore is already on at the beginning of the frame and turns off at some later time, or the fluorophore starts off and turns on during the frame. We neglect processes such as the fluorophore turning on then off and then back on within a frame as these will rarely occur, which is true if the exposure time is much smaller than the average OFF time, a condition well satisfied in SMLM experiments.

Now the probability of a fluorophore being ON at the beginning of a frame, at some time *t*, is given by *p*_*on*_(*t*). Likewise, the probability that a fluorophore is OFF at the beginning of a frame but turns ON at some point during the exposure time *τ*_*frame*_ is simply 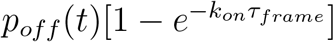. The discretized probability of finding a dye ON is then given by

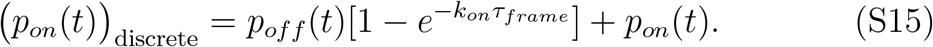

For the two-state model in the steady-state

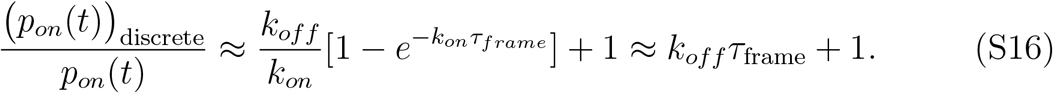

This last relation is justified since *k*_*on*_*τ*_*frame*_ *<* 10^−3^ in most SMLM acquisitions.

For the three-state model, *p*_*on*_(*t*) and *p*_*off*_ (*t*) may be approximated by the relations in Equations (S11) and (S13), respectively. Substituting these expressions in Equation (S15) yields

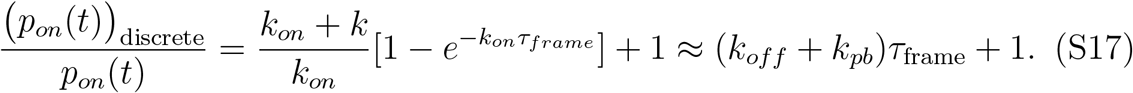

The effects of this scaling on the relative number of missed localizations is shown in Figure S2 for the case of the three-state photoswitching model with *k*_*off*_ = 0.01 ms^−1^, *DC* = 5 × 10^−3^, and *k*_*pb*_ = 1.25 × 10^−3^ ms^−1^ for three different frame lenghts. The relative number of missed localizations in the continuous case is also shown for reference. The scaling indicates that the frame size should be taken as small as possible to minimize the effects of discretization on increaing the probability of overlapping fluorophores. The lower bound for frame size would then be dictated by a high enough signal-to-noise ratio to minimize the number of localizations/blinks masked by the background. For imaging applications, a high localization precision may also be desired, which sets a second constraint on the smallest frame size possible. Both of these conditions can be met with fluorophores with a high photon emission rate.

**Figure S2:**
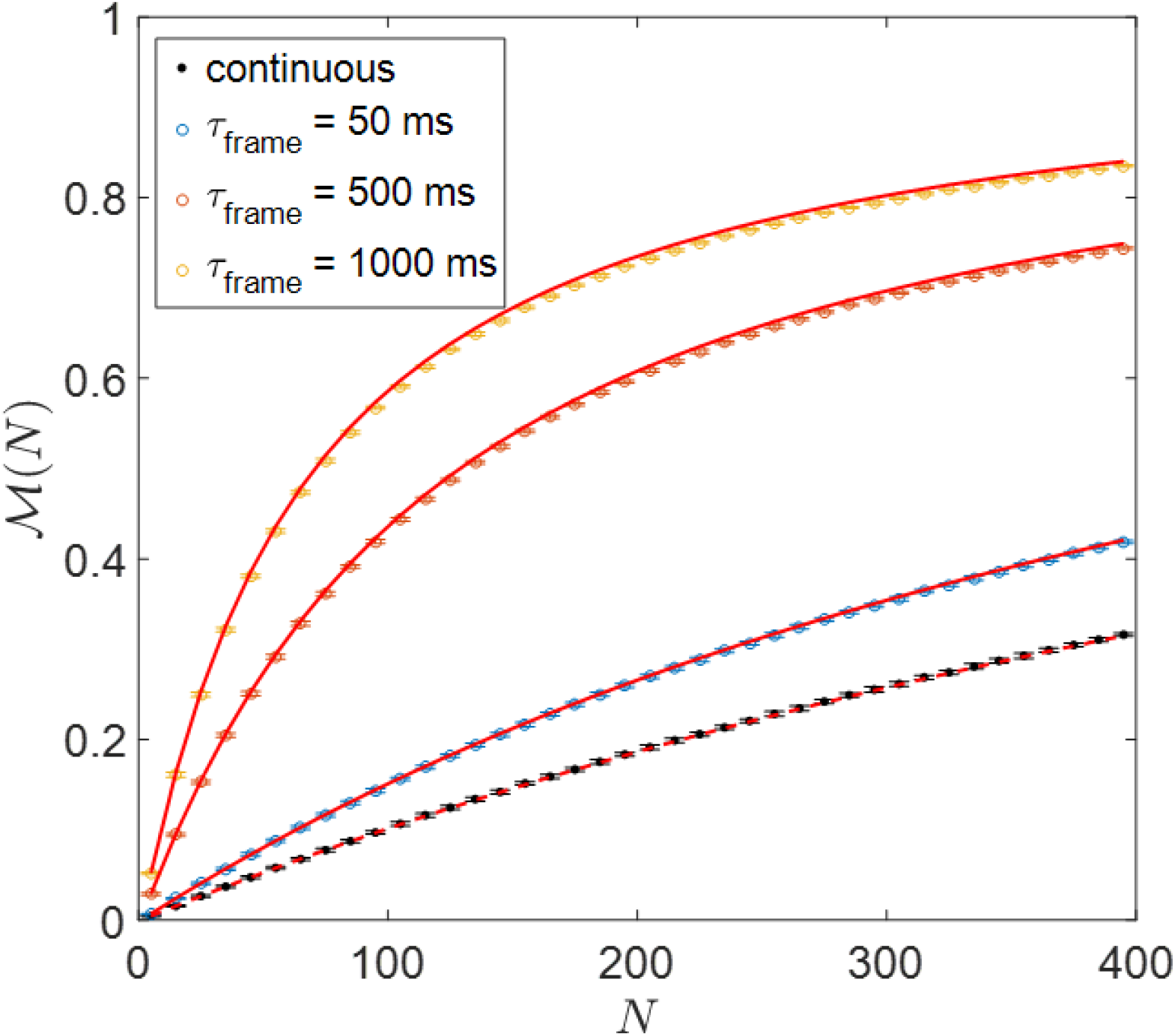
Discretization scaling shown for three different frame lengths (simulated data): 50 ms, 500 ms, and 1000 ms for photoswitching dyes with photobleaching (*k*_*off*_ = 0.01 ms^−1^, *DC* = 5 × 10^−3^, and *k*_*pb*_ = 1.25 × 10^−3^ ms^−1^). The solid (red) lines show the scaled theoretical estimate for the relative number of missed localizations with *τ*_*frame*_ = 50 ms, 500 ms, and 1000 ms in Eq. S17. The relative number of missed localizations is also shown for the continuous case, along with the theoretical estimate (dashed (red) line).

### 6 Off time distribution for Alexa647

**Figure S3:**
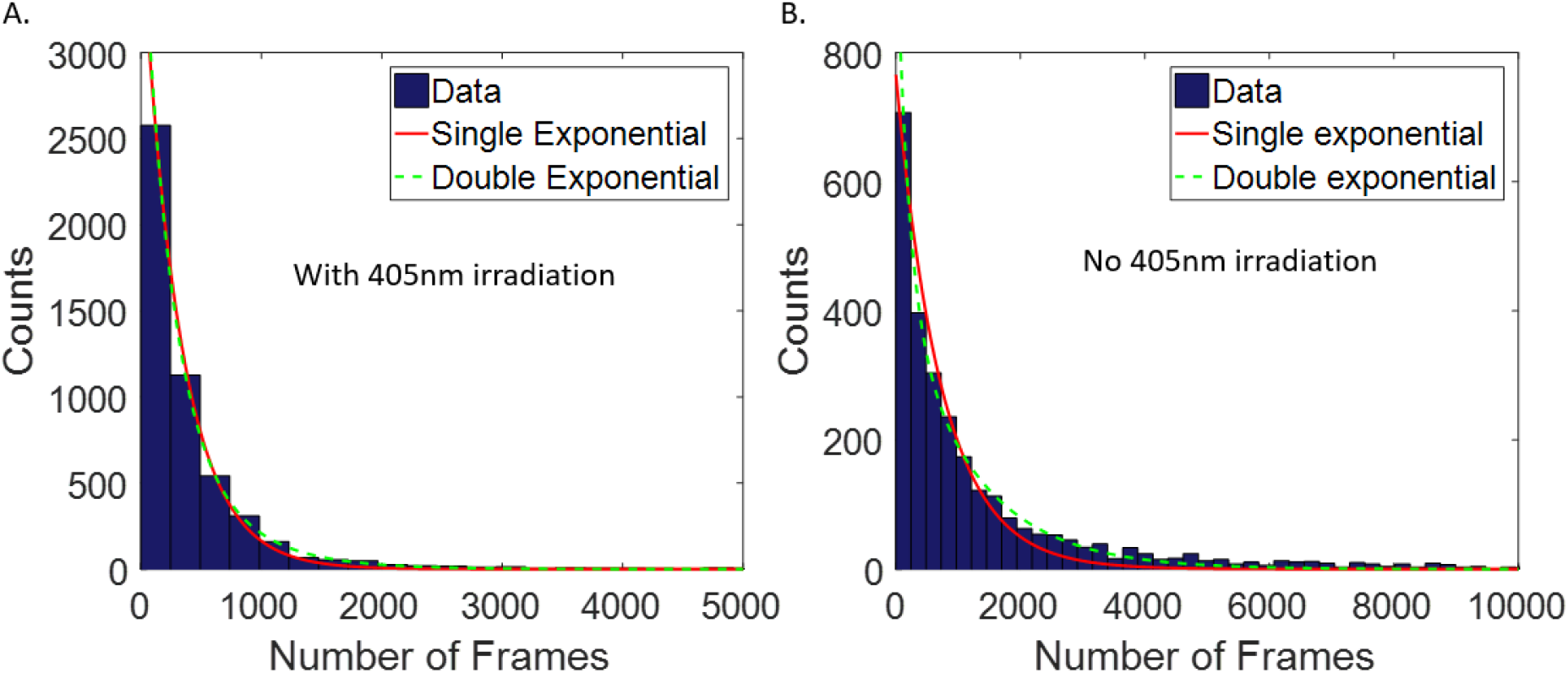
(A) Off time distribution for Alexa647 with 405 nm irradiation to induce recovery from OFF state. (B) Off time distribution for Alexa647 without 405 nm irradiation. Solid red line shows the fit to a single exponential model. Dashed green line shows the fit to a double model exponential. The presence of 405 nm irradiation appears to suppress the long tail contributions of the second exponential term, likely indicative of a second dark state.

### 7 Extrapolating from missed localizations to missed blinks

A number of SMLM-based counting techniques rely on counting the number of blinks per diffraction-limited spot to infer the total number of molecules. Estimating the fluorophore density for a given relative number of missed blinks is therefore also of interest to the design of counting experiments.

We can estimate a theoretical lower bound on the relative number of missed blinks for low fluorophore densities by the following argument. As done in the main text, we discretize time into a series of discrete bins of length *δt*. Suppose we now have a blink at some point in time that lasts for some average time *τ*_*on*_ (Fig. S4). If a second dye turns on at any of the bins indicated by the red arrows in Fig. S4, the two overlapping blinks will be counted as one blink. The probability that a second *blink* occurs at any of the indicated bins is simply *p*_*on*_(*t*)*/τ*_*on*_. So the probability of overlap scales like 2 × *p*_*on*_ and we pick up an additional factor of 2. Note, there is an additional bin in the illustration, making the factor slightly greater than 2, but for realistic ON times, this becomes negligible.

**Figure S4:**
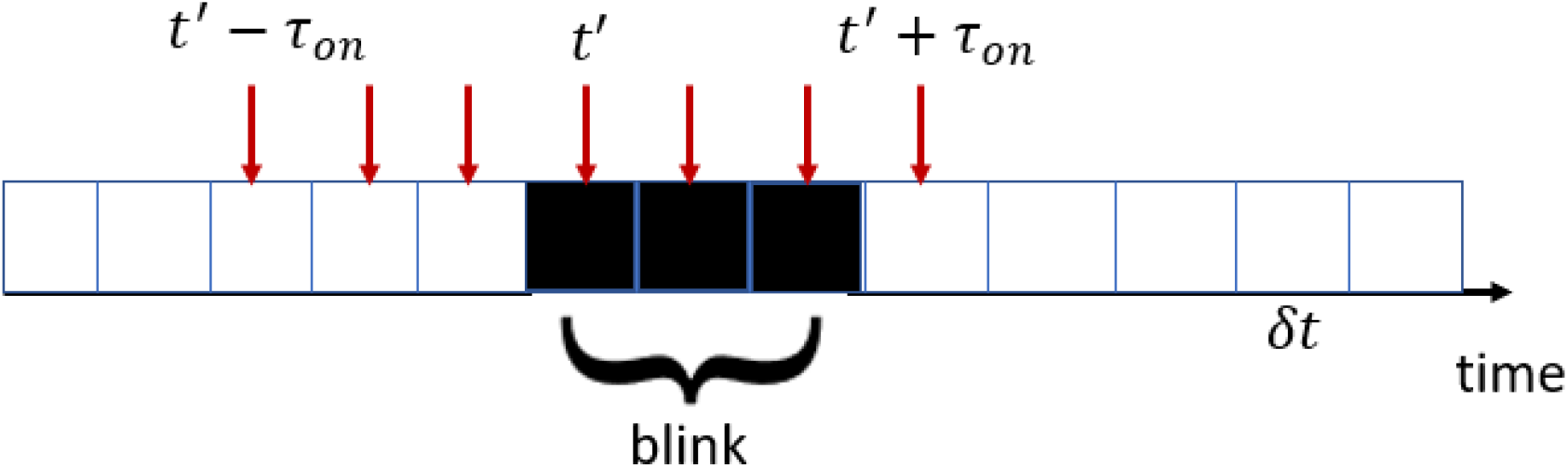
Representation of a dye blinking and the probability of being unable to distinguish it from another dye blinking. A dye turns on at time *t*′ and remains on for some time *τ*_*on*_ (measured in units of bin sizes of length *δt*) on average (shown here for *τ*_*on*_ = 3*δt*). A dye that turns on at any bin between *t*′ − *τ*_*on*_ and *t*′ + *τ*_*on*_ will result in overlapping blinks.

